# Ocelli: an open-source tool for the visualization of developmental multimodal single-cell data

**DOI:** 10.1101/2023.10.05.561074

**Authors:** Piotr Rutkowski, Marcin Tabaka

## Abstract

The recent expansion of single-cell technologies has enabled simultaneous genome-wide measurements of multiple modalities in the same single cell. The potential to jointly profile such modalities as gene expression, chromatin accessibility, protein epitopes, or multiple histone modifications at single-cell resolution represents a compelling opportunity to study developmental processes at multiple layers of gene regulation. Here, we present Ocelli, a lightweight Python package for scalable visualization and exploration of developmental multimodal single-cell data. The core functionality of Ocelli focuses on diffusion-based modeling of developmental processes. Ocelli addresses common tasks in developmental single-cell data analysis, such as visualization of cells on a low-dimensional embedding that preserves the continuity of the developmental progression of cells, identification of rare and transient cell states, integration with trajectory inference algorithms, and imputation of undetected feature counts. Extensive benchmarking shows that Ocelli outperforms existing methods regarding computational time and quality of the reconstructed low-dimensional representation of developmental data.

## Introduction

Multimodal data comprises distinct feature sets (called modalities or views) acquired by different detectors describing the same object [1, 2]. For example, a video includes modalities such as a video recording, an audio track, and usually text captions. Genome-wide single-cell technologies [3] can nowadays simultaneously measure multiple modalities from the same cell, including (i) transcriptome and chromatin accessibility [4–7]; (ii) transcriptome and DNA methylation [8]; (iii) transcriptome and chromatin proteins or histone modifications [9, 10]; (iv) transcriptome and protein epitope [11]; (v) transcriptome, chromatin accessibility, and protein epitope [12]; (vi) multiple histone modifications and DNA-bound proteins [13–17]; and (vii) open and closed chromatin [18].

Each single-cell data modality is high-dimensional. The number of measured features spans from hundreds in the case of protein epitopes to hundreds of thousands for chromatin-accessible sites. The multiple modalities profiled from the same single cell correspond to consecutive stages of gene expression, from its regulation by modifying chromatin architecture and engaging transcription-initiation proteins to the synthesis of mRNA and protein molecules. For example, chromatin becomes accessible before gene expression commences, resulting in temporal coherence between modalities. Thus, all modalities need to be modeled simultaneously to visualize the developmental progression or activation of cells and lineage branching accurately.

The inherent feature of data generated by single-cell technologies is a high level of technical noise resulting in data sparsity due to under-sampling of features from the same cell [19]. Additionally, different modalities have profoundly distinct statistical properties, which makes it challenging to find a joint representative data manifold. Modeling developmental processes is an additional challenge as cellular differentiation is a continuous and nonlinear process [20]. The origin of nonlinearity stems from the regulation of gene expression, where even the most straightforward regulation by a single transcription factor in an open-loop scenario results in nonlinear transcriptional kinetics that generates non-trivial gene expression patterns [21].

Multiple methods have been developed to deal with high-throughput high-dimensional unimodal single-cell data to study developmental processes. For example, Monocle [22] leverages a cellular minimum spanning tree (or DDRTree [23] in Monocle 2 and 3 [24]) to find dimensionally reduced gene expression space; Velocyto [25] and scVelo [26] model cellular transitions from transcriptional kinetics (named RNA velocity), leveraging the relationship between unspliced and spliced mRNA molecules within cells; Waddington-OT [27] employs optimal transport theory to find temporal couplings of differentiating cells; diffusion maps [20, 28–31] embed cells onto a low-dimensional manifold by eigendecomposing the diffusion operator. The significant advantages of diffusion maps are their robustness to noise, high sensitivity to finding rare and transient cell states [27], and ability to preserve nonlinear local and global structures, together with a continuum of developmental cell transitions. The structure and continuum are preserved because diffusion maps utilize a distance metric (diffusion time) relevant to differentiation processes [20, 30].

Recently, several methods have been developed to generate low-dimensional joint embeddings from multimodal single-cell data. The methods can generally be divided into deep learning and probabilistic categories. Deep learning [32] is the dominant branch of machine learning that gained popularity due to the increasing computational capabilities of modern computers [33]. While the variety of deep learning models is enormous, autoencoder architecture [34] has become the leading tool for visualizing multimodal single-cell data [35–38]. Autoencoder is a neural network that encodes and compresses complex, high-dimensional data. Its objective is to learn and perform two functions: an encoder that transforms the input data and a decoder that reconstructs the input data from the encoded representation. The performance of an autoencoder is measured by comparing input and reconstructed data - with the goal of them being similar. Autoencoders proved to be a potent model architecture for problems such as dimension reduction [32], drug discovery [39], and image processing [40, 41]. The advantage of deep learning methods is their theoretical ability to learn very complex functions [42]. However, in practice, they are prone to underfitting or overfitting. Additionally, they are computationally expensive and require extensive data sizes to converge. Probabilistic methods for visualizing multimodal single-cell data include Weighted Nearest Neighbors [43] (WNN), and Multi-Omics Factor Analysis v2 [44] (MOFA+). WNN integrates modalities by constructing a weighted nearest-neighbor graph based on a weighted average of modality similarities. The graph is then visualized with UMAP [45]. MOFA+ factorizes multimodal single-cell data to latent factors and corresponding feature weights. It uses stochastic variational inference [46, 47] for scalability of the parameter inference. Other approaches based on integrative methods exist, such as MOJITOO [48] that uses canonical correlation analysis to find low-dimensional multimodal data representation.

Here, we introduce Ocelli, a multi-part diffusion-based analysis and visualization strategy for developmental multimodal data. Ocelli leverages topic modeling and multimodal diffusion maps to reduce dimensionality and to preserve ordering, continuity, and branching of developmental trajectories. Explicit incorporation of modality-specific transition probabilities allows us to generate more informative developmental data visualization. Moreover, multimodal diffusion-based modeling provides an efficient solution to deal with such caveats of single-cell genomics data as sparsity of measured features. In addition, we show that Ocelli facilitates extraction and analysis of developmental subtrajectories from single-cell atlases. Ocelli was designed to work on an arbitrary number of modalities and optimized to handle scalability of current single-cell profiling technologies. Ocelli is available as an open-source software package at https://github.com/TabakaLab/ocelli along with tutorials and documentation at https://ocelli.readthedocs.io.

## Results

### Overview of Ocelli

Ocelli is an explainable multimodal framework (Fig. 1a and Methods) to learn a low-dimensional representation of developmental trajectories. In the data preprocessing step, we find modality-specific programs with topic modeling using Latent Dirichlet Allocation (LDA). This step reduces the dimensionality of feature space and denoises each modality while preserving the continuity of the data. Then, we model each modality as a diffusion process comprising affinities between neighboring cells.

**Fig. 1.**
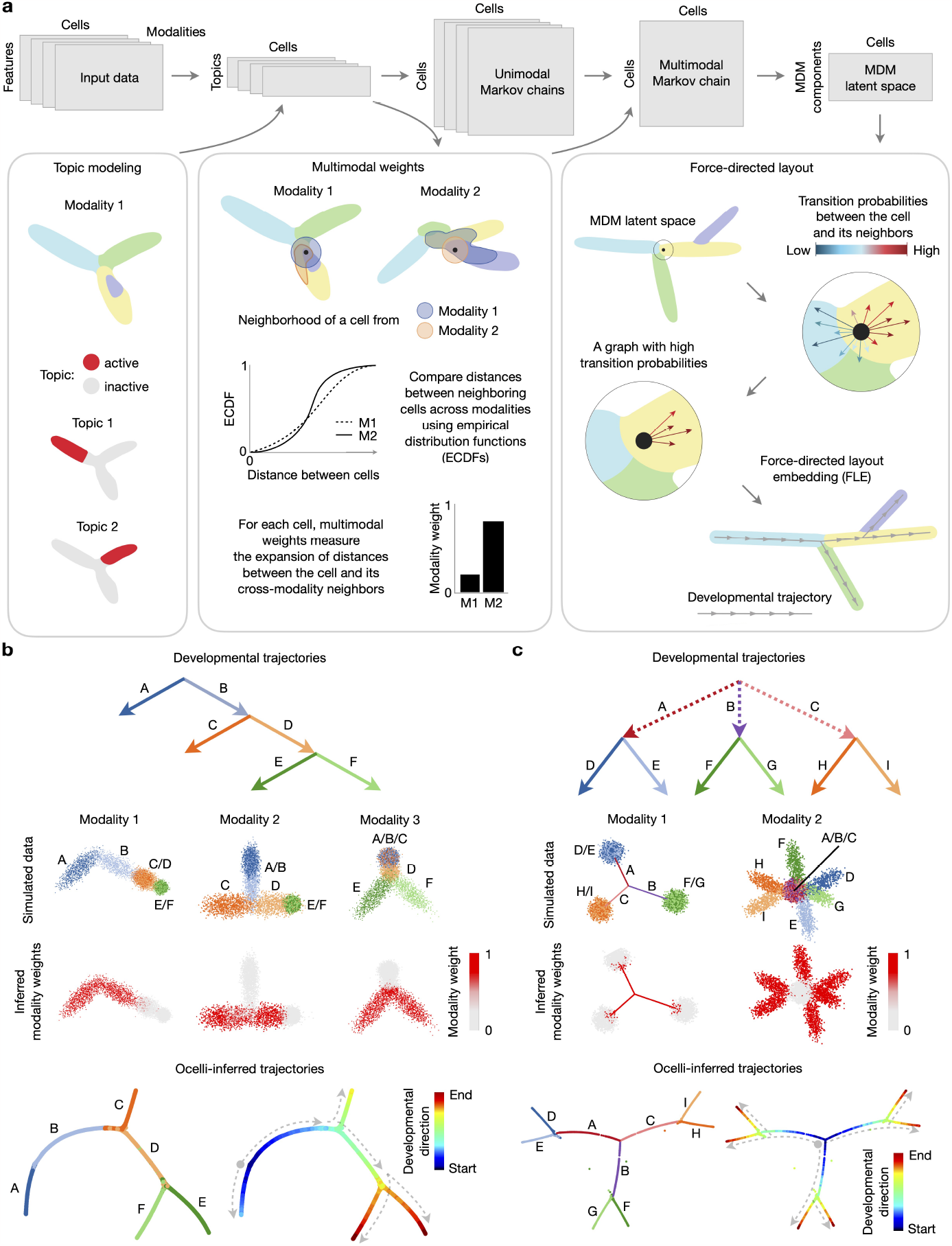
Ocelli overview. **a**, Schematic view of Ocelli workflow. The Ocelli core comprises multimodal dimension reduction with multimodal diffusion maps (MDM) and data visualization functionalities. Preprocessed modalities, modeled individually as diffusion processes, are merged into a weighted multimodal Markov chain and then decomposed into the MDM latent space. Ocelli can visualize MDM embeddings using a force-directed layout that employs cellular transition probabilities or any other dimension reduction algorithm, e.g., UMAP. **b-c**, Application of Ocelli to simulated data: the binary tree dataset with three modalities and six lineages A-F (**b**) and the rare transitions dataset with two modalities and nine lineages A-I (**c**). From top: a diagram of lineage trajectories (top), projections of modalities with lineage annotations and distribution of inferred multimodal weights (middle), an Ocelli embedding visualized using ForceAtlas2 with lineage annotations, and a developmental pseudotime progression (bottom).

In the second part of the model, we compute cell-specific modality weights that determine the importance of each modality for each cell from the topic latent representations. This approach resembles the WNN method. However, we compute cell-cell affinities using empirical cumulative distribution functions (ECDFs) of distances between cells. Their values encode distances with respect to the global structure of the topic latent space. Having multimodal weights, we construct the Multimodal Markov Chain (MMC) as a weighted sum of the unimodal affinities between cells. Multimodal weights quantify how informative modalities’ affinities are about the underlying developmental process. Next, we determine the latent space of multimodal diffusion maps (MDM) by factoring the MMC into eigenvectors and eigenvalues. This step further reduces the dimensionality of features, denoises the data, and allows us to find underlying developmental processes and cell ordering from the data. Then, we build the nearest neighbor cell-cell graph in the space of MDM components.

Finally, to obtain a latent space that reflects all modalities and additional information about cell fates, we leverage modality-inferred or -measured transition probabilities between cells to correct the cell-cell graph created from MDM. The transition probabilities between cells can be inferred from, e.g., RNA velocity [25, 26], optimal transport [27] algorithm, or determined from lineage-tracing experiments [49]. We modify the MDM nearest neighbors graph by rewiring edges to connect cells with the highest transition probabilities (see Methods for details). This step is used only for cells with determined transitions. The corrected cell-cell graph is visualized in 2D or 3D as a Force-directed Layout Embedding (FLE) [27] with ForceAtlas2 [50, 51] or Uniform Manifold Approximation and Projection (UMAP) [45].

### Modeling differentiation with Ocelli

First, we validated the analysis strategy of Ocelli in inferring developmental progression and lineage branching of cells using simulated data. We modeled the developmental process as a binary tree composed of three modalities with two lineages branching in each modality (Fig. 1b). Ocelli identified the importance of each modality, reconstructed the branching points, and preserved the continuity of developmental trajectories with high accuracy. Next, we considered a developmental process with transient developmental transitions in the first modality (Fig. 1c, lineages A-C) followed by lineage branching in the second modality (lineages D-I). Ocelli displayed remarkable sensitivity in reconstructing transient developmental transitions and lineage branching (Fig. 1c). Finally, we compared the visualizations generated from simulated multimodal data between Ocelli and WNN, a direct competitor graph-based method (Supplementary Fig. S1a-c). Ocelli generally generates more coherent and compact visualizations of trajectories while preserving their continuity and recovering lineage branching (Supplementary Fig. S1a). Comparison of distributions of modality weights between the methods (Supplementary Fig. S1b) shows that Ocelli accurately infers the distributions (Supplementary Fig. S1c), minimizing the mean squared errors between ground truth and inferred modality weights. This is most apparent when comparing inferred modalities and generated visualizations for transient trajectories (Supplementary Fig. S1a-c, right panel).

Having demonstrated its performance on simulated data, we then used Ocelli to infer lineage dynamics of the regenerative subset of the hair follicle (Fig. 2a) from a dataset generated from simultaneous profiling of chromatin accessibility and gene expression by the SHARE-seq method [6]. Ocelli deconvoluted lineages building the hair shaft (HS) and inner root sheath (IRS) from transit-amplifying cells (TACs) (Fig. 2b), correctly identifying the higher weighting of chromatin accessibility in progenitor cells (TACs) and of RNA in differentiated cells (Fig. 2c,d). The obtained visualization shows two populations of progenitor cells and four populations of terminal differentiated cells. To annotate these cell populations, we computed gene signature activities determined from Smart-seq2 dataset [52]. Ocelli clearly distinguished hair shaft cells (medulla (HS-Me), cortex (HS-Co)) and inner root sheath cells (Huxley’s layer (IRS-Hu), Henle’s layer (IRS-He)) (Fig. 2e). In addition, CellRank [53] analysis confirms four terminally differentiated states that overlap with these gene signature activities (Fig. 2f).

**Fig. 2.**
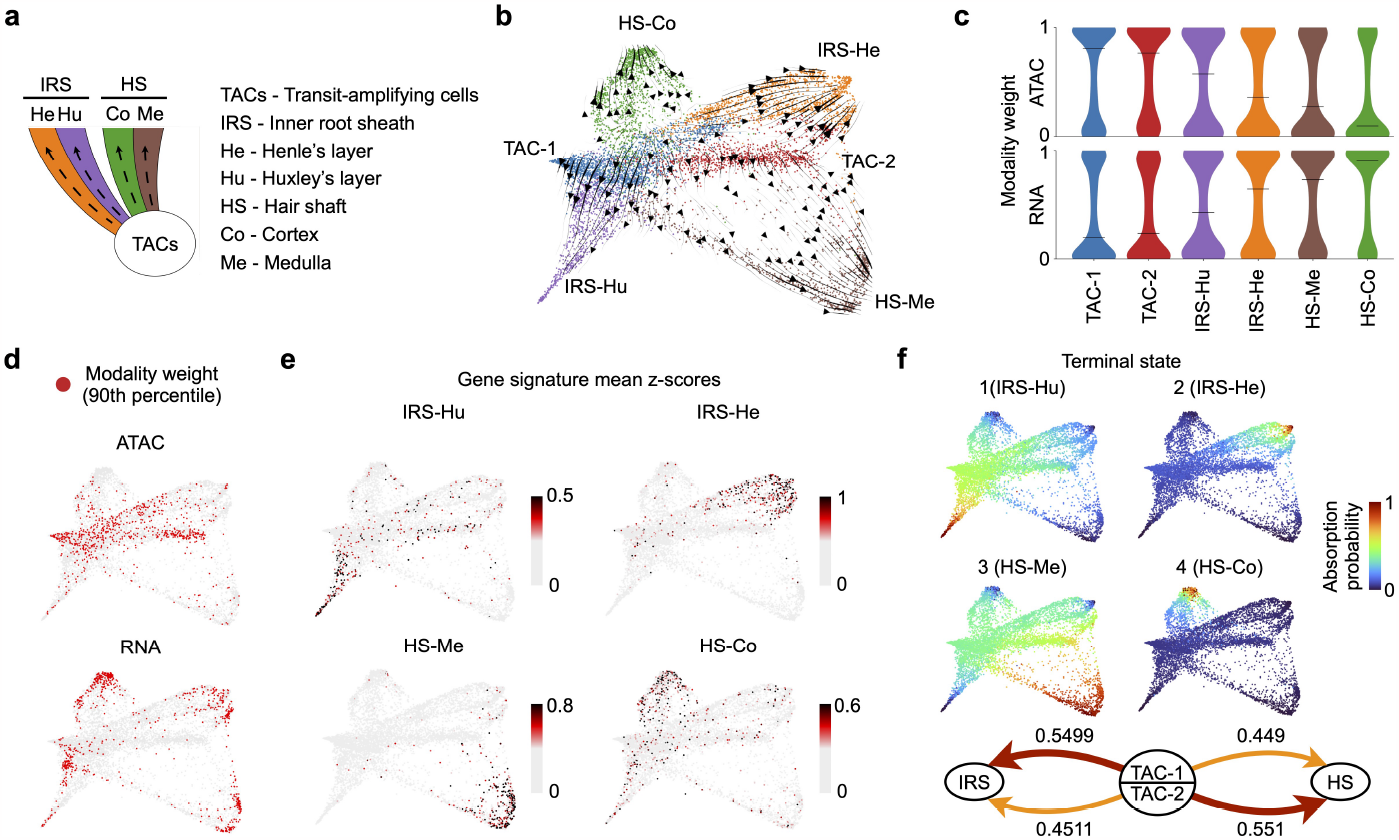
Reconstruction of developmental pathways in the hair follicle SHARE-seq dataset, comprising chromatin accessibility and transcriptome modalities. **a**, Schematic depicting the differentiation pathways in the regenerative part of hair follicle. **b**, Ocelli’s visualization of the single-cell SHARE-seq data of the regenerative part of hair follicle. **c**, Inferred chromatin accessibility weights show higher levels for progenitor transit-amplifying cells (TACs). In contrast, the transcriptome weights are elevated for differentiated cells. **d**, A distribution of 10 % of cells with the highest weights of each modality. **e**, Gene signature activity [52], computed as gene expression mean z-scores, of hair shaft: medulla (HS-Me) and cortex (HS-Co); and inner root sheath: Huxley’s layer (IRS-Hu) and Henle’s layer (IRS-He). **f**, CellRank [53] analysis detects four terminal differentiation states. Mean absorption probability shows a variety within transit-amplifying cells (TAC). Inner root sheath (IRS) cells are more likely to develop from TAC-1 and hair shaft (HS) cells from TAC-2.

### Benchmarking analysis

We further performed extensive benchmarking of Ocelli against other recently developed methods for single-cell multimodal data analysis and visualization regarding computational runtime and quality of the generated data embeddings. For the latter benchmarking analyses, we wanted to explore the quality of the visualization of developmental single-cell data focusing on the regenerative part of hair follicle (Fig. 2). While some benchmarking strategies have been developed for assessing analysis and visualization of well-clustered data, such as silhouette score or adjusted Rand index score [48], we designed new benchmarks related to the visualization of developmental multimodal data and transitions between cells (Fig. 3, Supplementary Fig. S2, Methods). Ocelli was benchmarked against (i) graph-based method, WNN [43]; (ii) statistical framework for multimodal factor analysis, MOFA+ [44]; (iii) deep learning-based methods - Cobolt [35], scMM [36], Matilda [37], MultiVI [38]; and (iv) canonical correlation analysis method MOJITOO [48] (Supplementary Fig. S2a, Methods).

**Fig. 3.**
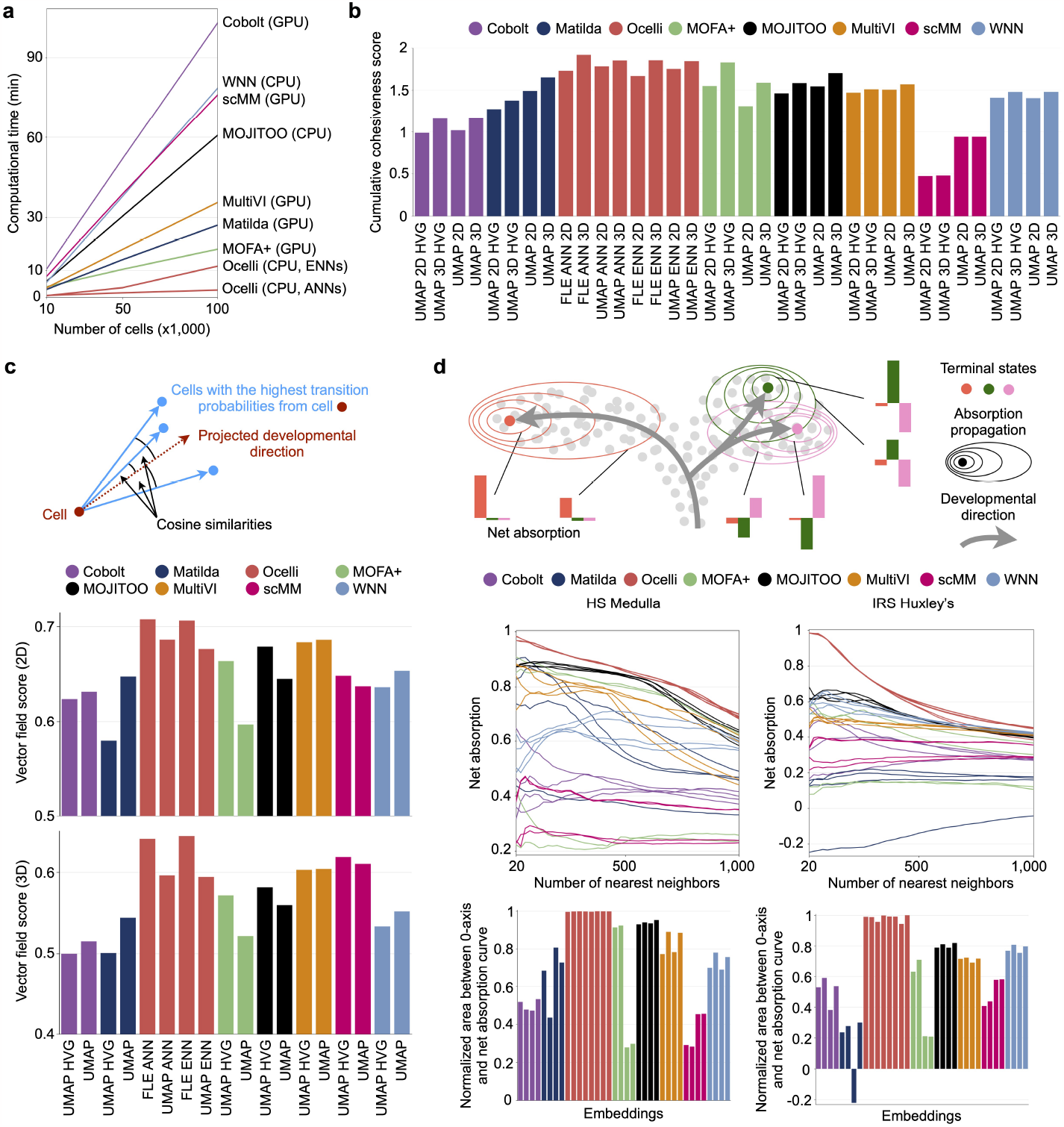
Benchmarking Ocelli against competing multimodal dimensionality reduction methods on a SHARE-seq dataset of the regenerative part of hair follicle. **a**, Ocelli improves computation time compared to state-of-the-art methods. **b**, The cohesiveness score quantifies the tendency of grouping cells with similar transcriptomic and chromatin accessibility signatures. MDM embeddings were computed for exact (ENN) and approximate nearest neighbors (ANN). Embeddings computed on 1,000 highly variable genes are marked as HVG. **c**, The vector field score measures the alignment of an embedding and the RNA velocity vector field. The score is the median cosine similarity between the cell’s projected developmental direction and directions toward cells with the highest RNA velocity transitions. **d**, Net absorption evaluates the separation of developmental terminal states in embeddings. Net absorption of a terminal state is its mean neighborhood absorption after subtracting mean absorptions to remaining terminal states. High net absorption translates to a well-separated terminal state. Embeddings are ordered as in Fig. 3b.

First, Ocelli exhibits improved computational runtimes compared to other competing multimodal methods (Fig. 3a). For example, reconstructing a joint embedding of 100,000 cells took Ocelli less than 3 minutes, outperforming all competing multimodal methods. We used GPU-accelerated computations for MOFA+ and deep learning methods and CPUs for others for these speed tests. We excluded preprocessing steps and used the same data dimensions to perform fair comparisons between the methods (Methods). For Ocelli, we determined the runtimes for exact (ENN) and approximate (ANN) approaches of the computation of nearest neighbors.

Next, we assessed how the aggregate signal of related features in each modality is displaced on the embedding. We used gene signature scores for the RNA modality. In the case of the ATAC modality, we leveraged transcription factor motif scores (Methods). A high-quality, developmental data embedding should group cells of similar scores together. A failed embedding can have high signature scores, for example, related to the same cell identity separated, which would suggest alternative lineages. To quantify this, we introduced the signature cohesiveness score (Methods). The evaluated embedding is dissected into a connectivity graph using the DBSCAN [54] algorithm. Then, each connected graph component is scored based on the presence of the most signature-active cells and the signature activity in their neighborhoods. The cohesiveness score favors an embedding in which most signature-active cells are grouped in a single connected graph component and if neighborhoods of highly signature-active cells are also highly signature-active. We considered different scenarios for data visualization (Methods): (i) optimal preprocessing steps for each method suggested by developers of each method; (ii) usage of all features or initial reduction of data dimensionality by selecting highly-variable features; (iii) final data embedding on two-dimensional (2D) or in three-dimensional (3D) using UMAP or FLE. The cumulative cohesiveness score for RNA and ATAC modality is the highest for Ocelli’ MDM and 3D force-directed layout embedding (Fig. 3b). Ocelli generally generates the most robust visualizations in this benchmark across all methods.

Finally, we developed two benchmarks to compare the accuracy of visualizing developmental trajectories across the methods. The first benchmark quantifies the reconstruction of temporal cell ordering and differentiation direction on the embedding in relation to the inferred transitions by the RNA velocity-based method [26]. The output of the method, RNA velocity vector field, is obtained from relative abundances of immature (unspliced) and mature (spliced) RNA molecules. Usually, it is visualized on data embeddings by arrows, depicting the cell’s projected developmental direction. Descendant cells with the highest transition probabilities should be aligned near the projected developmental direction (Fig. 3c) in case of high-quality data embedding. Across all cells, we quantified this by computing cosine similarities between the projected developmental direction and the direction toward cells with the highest transition probabilities. 2D and 3D data embeddings generated by Ocelli using FLE show the highest scores in the ordering of cells according to RNA velocity vector fields across all tested methods (Fig. 3c). As a second quantitative benchmark, we determined the separability of cell lineages on generated data embeddings. Using CellRank’s [53] determined terminal states based on RNA velocity results, we tested how the cell neighborhoods of terminal states are displaced on the embedding relatively to each other (Fig. 3d). To do so, first, we determined cells’ absorption probabilities to terminal states using CellRank algorithm. Then, we systematically increased the number of nearest neighbors of one terminal state and computed, among the nearest neighbors, the difference between mean absorption probabilities to this terminal state and to others (Fig. 3d, Methods). This net absorption score is high if cell trajectories of different lineages are well separated on the embedding. We computed the net absorption scores to four terminal states of hair follicle for all tested methods (Fig. 3d, Supplementary Fig. S2b). Regardless of the final data embedding strategy (UMAP or FLE), Ocelli outperformed all other methods in separating trajectories to all terminal states.

### Diffusion-based multimodal imputation

The single-cell profiling of cells inevitably generates data of high sparsity levels, resulting in 0-inflated feature values that obstruct the inference of relationships between features of the same or different types. The data’s sparsity originates from inefficient feature detection in single cells ranging from 10-45% of expressed RNA molecules in scRNA-seq and 1-10% of accessible sites in scATAC-seq [55]. To address the sparsity observed in single-cell data, many unimodal imputation methods have been developed in recent years [56]. In general, the imputation methods can be categorized into three categories: (i) probabilistic ones that directly model the data sparsity; (ii) smooth-based methods that smooth or diffuse feature values among nearby cells; and (iii) deep learning methods that first identify latent spaces and then reconstruct the feature matrices. The extensive benchmarking [56] of scRNA-seq data imputation methods revealed that smooth-based approaches, e.g., MAGIC [19] and kNN-smoothing [57] are the top unimodal imputation methods. MAGIC uses the diffusion operator to first learn data manifold [28] and then restores the missing expression values by exponentiation of the Markov matrix to a power *t* and multiplying the result by original expression matrix. While the unimodal imputation methods can be bluntly applied to each modality separately, multimodal imputation has the potential to resolve more nuanced differences between cells along the differentiation axis and, thereby, improve imputations.

Here, we demonstrate several key properties of Multimodal Diffusion Maps to impute high-dimensional signals in multimodal data in a highly efficient way. First, instead of exponentiation of the multimodal Markov matrix, which is time-consuming, we eigendecompose it (Fig. 4a). Then, the exponentiation of the Markov matrix is equivalent to the exponentiation of eigenvalues, which is of constant complexity as the number of diffusion steps *t* varies. It results in highly scalable and efficient imputing of unobserved data for a large number of cells (Fig. 4b, upper panel) and features (Fig. 4b, lower panel). The salient feature of imputations is the recovery of gene-gene relationships and regulatory interactions [19]. First, we tested the imputations on two canonical markers of Henle’s lineage, Krt71 and Krt73. Ocelli’s multimodal imputations recovered the correlations between the expression of the marker genes (Fig. 4c) and the chromatin accessibilities at their promoters (Fig. 4d). To further test the usability of Ocelli’s multimodal imputations, we explored the recovery of correlations between features from different modalities or among cells. In the first case, we imputed gene expression, promoter, and distal element accessibility for 50 top gene markers of clusters identified earlier as terminal states. Ocelli restored the coactivity of regulatory elements and genes within differentiated cells (Fig. 4e). The second case involved imputing the dataset’s expression values of all genes and regulatory elements. Cell-to-cell correlation heatmaps of Ocelli-imputed RNA estimates (*t* = 1) (Fig. 4f) and Ocelli-imputed promoter and distal elements accessibility estimates (*t* = 1) (Fig. 4g) showed that multimodal imputations help to reconstruct similarities between closely related cells.

**Fig. 4.**
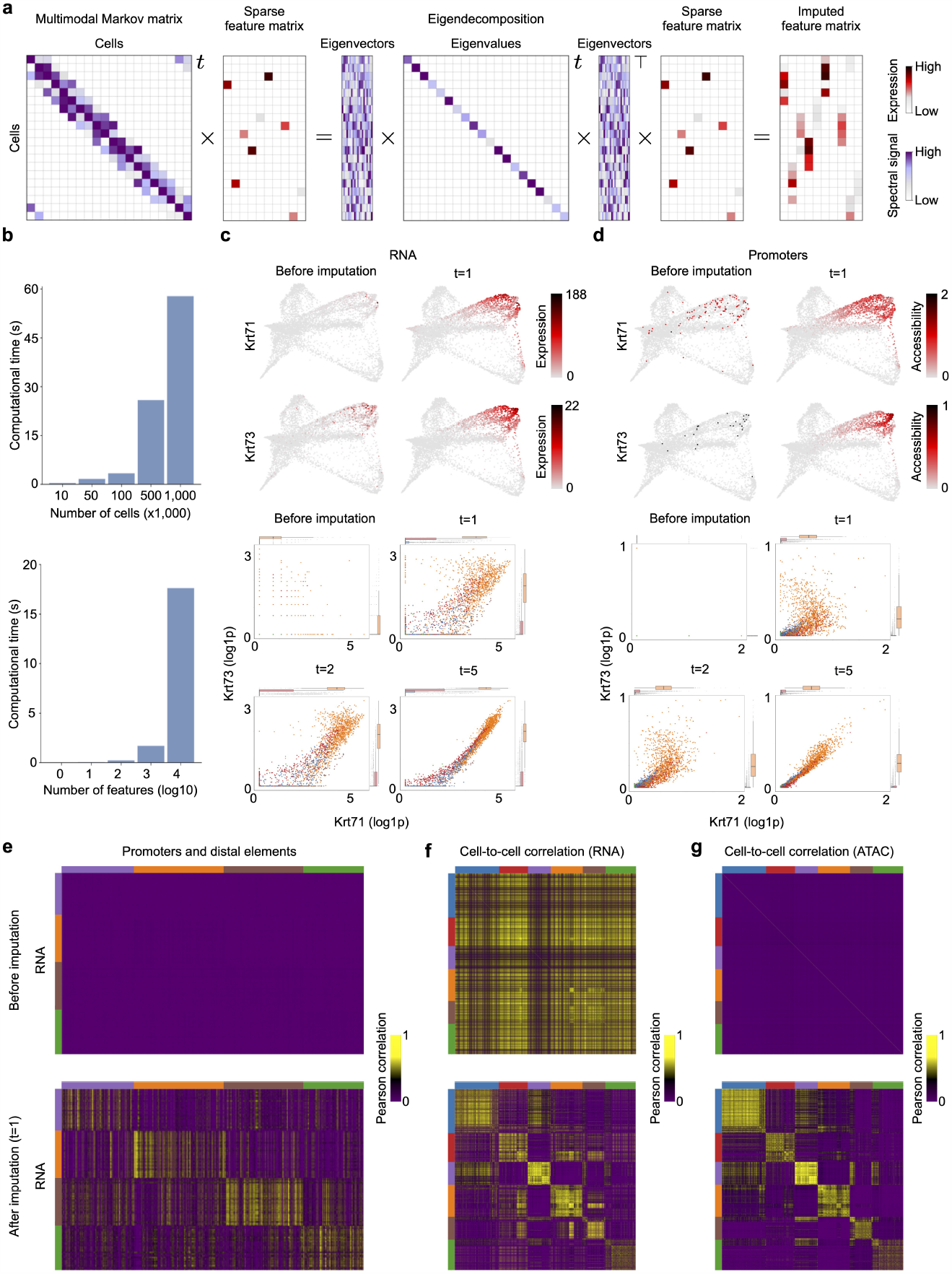
Ocelli enables the reconstruction of highly sparse features of each modality using multimodal imputations. **a**, The exponentiation of the multimodal affinity matrix is equivalent to eigendecomposition with exponentiated eigenvalues, significantly improving the computational time of imputations. **b**, The Ocelli’s runtimes for multimodal imputation for the increasing number of cells (top) and the number of features being imputed (bottom). **c**, Gene expression levels (top) and scatterplots of gene-gene relationships (bottom) for two canonical markers of the IRS Henle’s layer (Krt71 and Krt73) with different amounts of diffusion. **d**, Chromatin accessibility of promoters for the markers in (**c**). **e**, Correlations of gene expression to chromatin accessibility of promoters and distal elements before and after imputations (*t* = 1) with Ocelli for differentiated cells. **f, g**, Cell-to-cell correlations computed on gene expression (**f**) and chromatin accessibility (**g**) levels before and after imputations (*t* = 1) with Ocelli.

### Diffusion-based multimodal exploration of subtrajectories

Building developmental single-cell atlases involves refined analysis of developmental subtrajectories from data consisting of multiple distinct lineages and cell types unrelated directly to a subtrajectory of interest [58]. Currently, such analysis entails clustering all cells, manual annotation of cell clusters, and supervised extraction of cell clusters that may belong to the studied subtrajectory. Here we show that data diffusion can facilitate the analysis of subtrajectories by unsupervised extraction of phenotypically related cells.

To illustrate multimodal diffusion-based extraction and refined reconstruction of a subtrajectory, we analyzed ASAP-seq human native hematopoietic differentiation data [12], obtained from simultaneous measurement of chromatin accessibility and protein epitopes (Fig. 5a) for 10,927 human bone marrow cells. First, consistent with the analysis results for the hair follicle dataset, Ocelli reconstructed the hematopoietic cell lineages with high accuracy, correctly assigning the highest chromatin accessibility weights to Hematopoietic Stem Cells (HSCs), gradually decreasing with more differentiated states of cells for which protein epitope modality becomes more prevalent (Fig. 5b,c). The vast majority of cells present are CD14+ monocytes, B, and T cells, which obstructs visualization of the hematopoietic hierarchy of cells. We sought to refine the analysis of the HSCs differentiation trajectories. We selected a CD34+ positive cluster of 351 cells and assigned the uniform distribution equal to 1 for the selected cells and 0 otherwise (Fig. 5d). Using the same data diffusion strategy as in imputations (Fig. 4a), we performed diffusion to expand the initial cell set to similar cells along the multimodal affinity-based graph structure. Finally, for diffusion time *t* = 20, the expanded set of 1,451 cells has been used as an input in Ocelli for multimodal visualization of the cells. The continuous trajectories from HSCs are readily apparent in the refined visualization of cells (Fig. 5e) which accurately reconstructed the early hematopoietic hierarchy [59, 60]. Moreover, the refined visualization shows distinct pathways of basophilic/mast cell and B cell progenitors that were buried in a coarse visualization of human bone marrow cells (Fig. 5a).

**Fig. 5.**
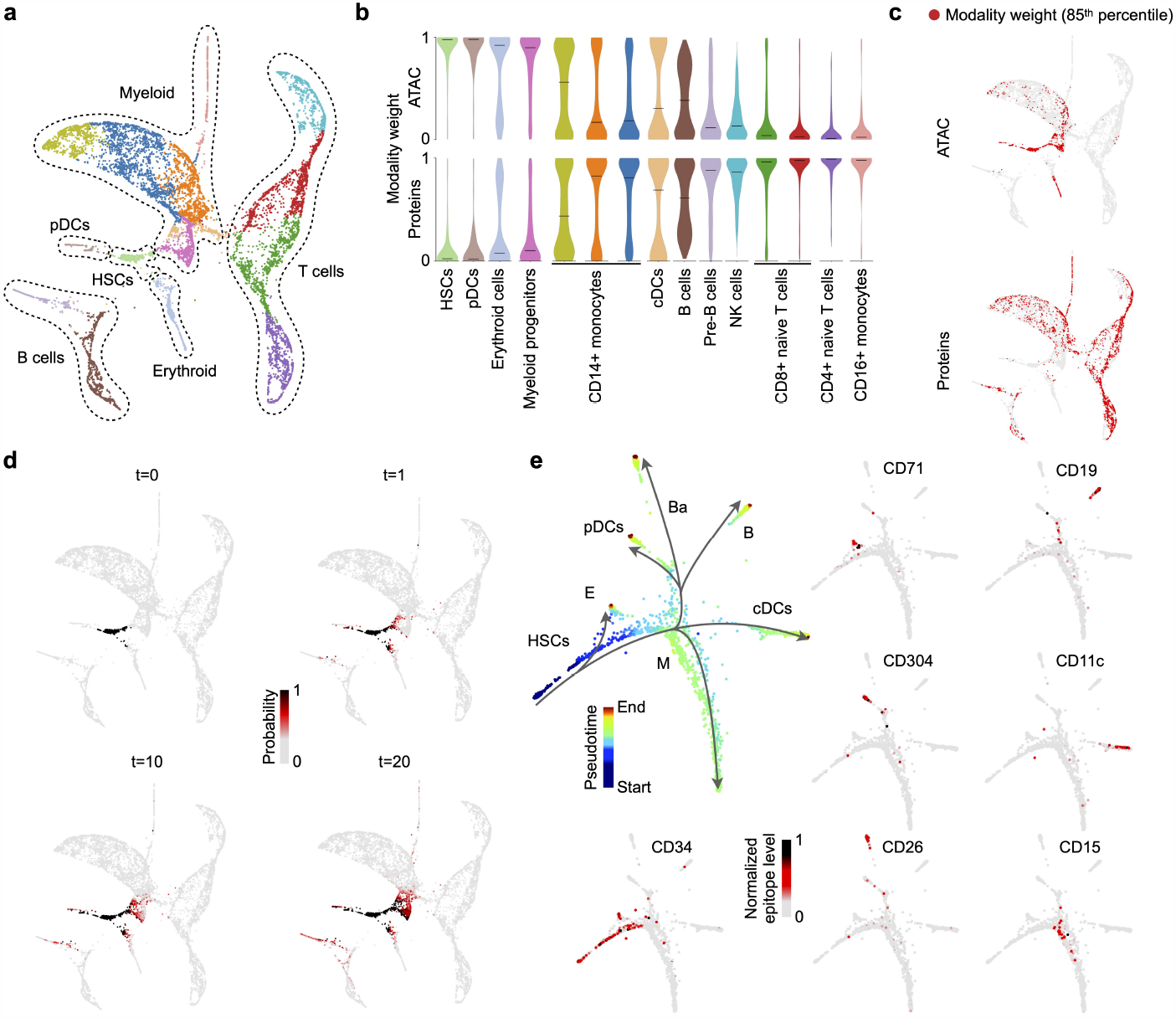
Cell selection strategy for refined analysis of trajectories. **a**, The multimodal visualization of ASAP-seq human bone marrow data with coarse annotation of hematopoietic cell lineages. **b**, Violin plots show a distribution of multimodal weights in each cluster from panel **a. c**, A distribution of 15 % of cells with the highest weights. **d**, Cells from the HSC cluster were assigned a uniform distribution equal to 1 (*t* = 0), and the diffusion was performed for different diffusion times *t* = 1, 10, 20. **e**, Cells selected at *t* = 20 were reanalyzed with Ocelli. The FLE plots show refined hematopoietic trajectories with cells colored by pseudotime or epitope level of lineage-specific protein markers. HSCs, hematopoietic stem cells; E, erythroid progenitors; pDCs plasmacytoid dendritic cells; Ba, basophilic/mast cell progenitors; B, B cell progenitors; cDCs, classical dendritic cells; M, myeloid progenitors.

### Utilizing Ocelli for single-modality data visualization

Gene expression programs (GEPs) determine cell identity and activity. GEPs are modules of coregulated genes [61–63]. GEPs maintain specific cell types and perform complex cellular activities such as proliferation, apoptosis, metabolism, differentiation, or responses to environmental cues. Each cell is a mixture of GEPs, and their relative contributions change continuously throughout cellular differentiation. Usually, GEPs are computationally determined in an unsupervised manner from scRNA-seq data by LDA [64] or non-negative matrix factorization [63]. We hypothesized that GEPs, inferred from unimodal scRNA-seq, could be leveraged directly as latent modalities to generate visualizations of scRNA-seq data with Ocelli. In this context, inference of GEPs in the first step would denoise single-cell data, and the multimodal analysis strategy implemented in Ocelli would accurately separate cell lineages and their states of different activities.

Here, we tested multimodal analysis framework on two developmental scRNA-seq datasets: (i) pancreatic endocrinogenesis [65], and (ii) cellular reprogramming to induced Pluripotent Stem Cells (iPSCs) [27]. First, we inferred GEPs using LDA analysis, where each LDA’s topic groups features into programs. Ocelli interprets each program as a modality, using the cell-topic distribution as modality weights. Then, we computed a low-dimensional MDM embedding on log-normalized topic-based latent modalities. Finally, we created a visualization of single cells with ForceAtlas2, leveraging inferred transition probabilities between cells. In the case of pancreatic endocrinogenesis data visualization, we used RNA velocity-derived transition probabilities. Ocelli accurately reconstructed complex developmental lineages (Fig. 6a), rare and transient cellular transitions (Fig. 6b), and different proliferative cell states (Fig. 6c-e). In detail, four hormone-producing endocrine cells, Alpha, Beta, Epsilon, and Delta, arise from the endocrine progenitor cells originating from cycling Ductal cells. The first progenitor branching point leads to the development of Epsilon cells and an intermediate cell population that generates independently Alpha cells [66]. Subsequently, progenitors further develop into Alpha, Beta, and Delta lineages. Delta cells are produced primarily after E15.5, with only a few cells detected at E15.5 [66]. Notably, Ocelli visualized (i) both differentiation pathways to Alpha cells: through intermediate cell population with Epsilon cells and then directly from endocrine progenitor cells; (ii) a distinct pathway to Delta cells despite its scarcity at the embryonic day E15.5; and (iii) a cell cycle of Ductal cells. Ocelli’s visualization shows concise trajectories as compared to competitive approaches [26].

**Fig. 6.**
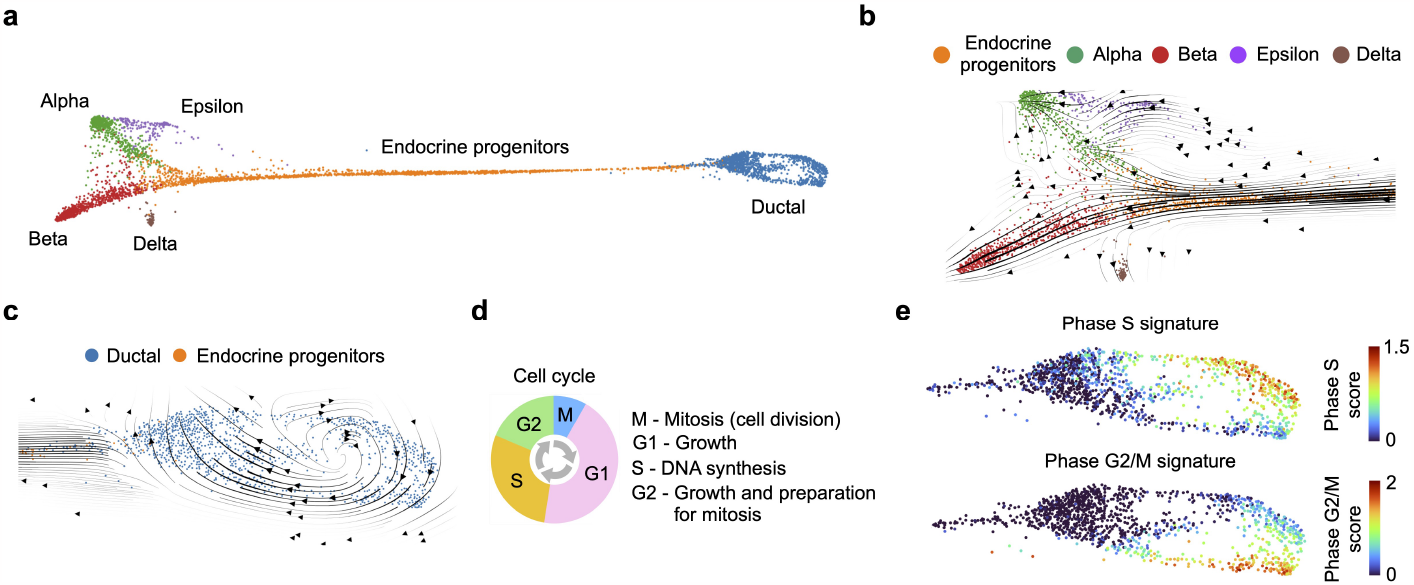
Exploration of the pancreatic endocrinogenesis unimodal dataset with Ocelli. **a**, The visualization of the cell trajectories generated by Ocelli. **b**, scVelo’s RNA velocity vector field projected onto developmental trajectories of endocrine cells reconstructed with Ocelli. **c-e**, Reconstruction of the cell cycle of Ductal cells. **c**, scVelo’s RNA velocity vector field showing inferred cell cycle trajectory of Ductal cells. **d**, A schematic view of cell cycle phases. **e**, Scores of cell cycle phases calculated using the list of cell cycle signature genes [84].

In the second example, reprogramming of cells to iPSCs, we leveraged Waddington-OT inferred cell-cell transition probabilities. The transition probabilities were obtained by the optimal transport modeling of the cell fates between two consecutive time points at which the cells were collected for single-cell profiling. Similarly to previous examples, Ocelli showed high sensitivity to capture precisely developmental branching points and to reconstruct rare cell transitions (Fig. 7). For example, Ocelli visualized a transient developmental trajectory to induced pluripotent stem cells (iPSCs) in serum condition (Fig. 7a,b), comprising as little as 0.45% of all cells at day 11. In addition to all previously annotated cell types [27], Ocelli was able to visualize a distinct population of keratinocytes depicting an asynchronous multiple-stage interfollicular differentiation [67] (Fig. 7c). Moreover, rare populations of extraembryonic endoderm (XEN) and trophoblast-like cells with their differentiation trajectories are clearly distinguishable (Fig. 7d-g). Earlier analyses [27] could not capture such precise trajectories and cell subtypes in a single, joint visualization. Taken together, Ocelli produces highly sensitive and accurate embeddings capable of showing rare cell transitions and multiple differentiation trajectories.

**Fig. 7.**
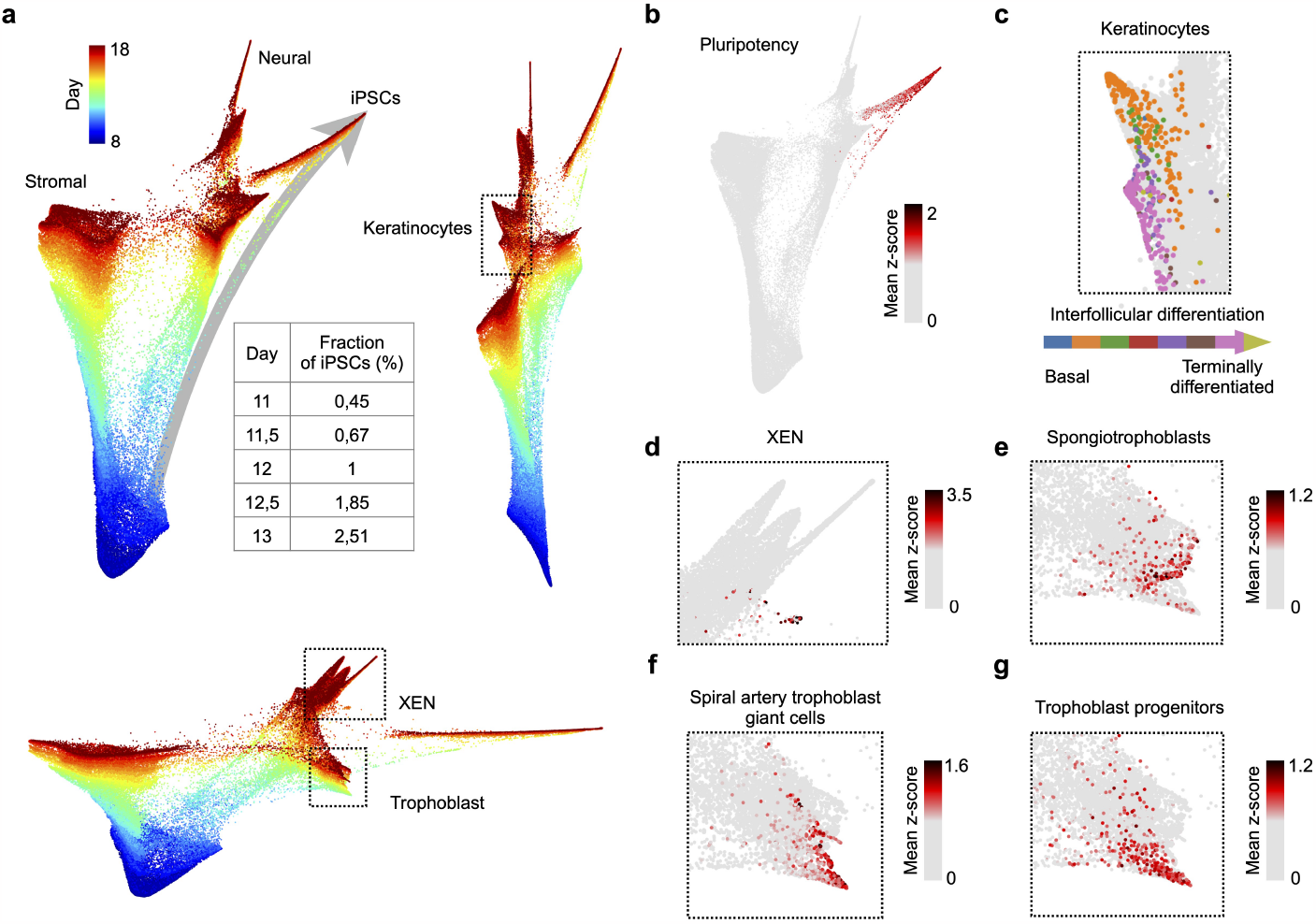
Sensitivity of Ocelli in reconstructing rare cell transitions. **a**, Reconstruction of cell trajectories in serum from the cell reprogramming dataset [27]. Three scatter plots show projections of a 3-dimensional embedding constructed with Ocelli. The arrow highlights a rare developmental trajectory of induced pluripotent stem cells (iPSCs). The table specifies the fraction of iPSCs concerning all cells from a given timestamp. **b-g**, Ocelli captures multiple developmental trajectories on a single joint embedding. Plots show the activity of gene signatures related to cell identities [27]: pluripotency (**b**); keratinocytes at differentiation stages from basal to terminally differentiated cells, where each cell is colored by its dominant differentiation phase based on the highest z-score (**c**); extraembryonic endoderm stem (XEN) cells (**d**); spongiotrophoblasts (**e**); spiral artery trophoblast giant cells (**f**); trophoblast progenitor cells (**g**).

## Discussion

Ocelli is an open-source visualization framework that uses multimodal diffusion maps to enhance multimodal single-cell sequencing data analysis and exploration. The framework was specifically designed to resolve developmental processes and diminish caveats of single-cell genomics data, such as data sparsity. Ocelli generates informative visualizations of developmental multimodal data and outperforms state-of-the-art multimodal algorithms regarding the computational runtime and the quality of reconstructed developmental trajectories. To our knowledge, Ocelli is the only method that learns low-dimensional embedding of the data coalescing information from both highly-dimensional multimodal features and modality-specific transition probabilities between cells. As a diffusion-based method, Ocelli enables scalable multimodal imputations of missing feature activity levels and has the potential to facilitate the exploration of trajectories from large datasets e.g. mammalian organogenesis [58], which typically consist of many unrelated differentiation subtrajectories or in which some of the ancestral states are missing. Finally, Ocelli can be exploited on unimodal single-cell (e.g., scRNA-seq) data to generate more informative visualizations with the prospective to better resolve cell identities, activities, and spare transitions along cell’s differentiation trajectories.

The ability of multimodal algorithms to deconvolute and interpret multi-faceted data, termed explainability, is one of the critical factors that govern their applicability to resolve biological processes. In other words, a method producing highly interpretable results can be preferred over a slightly more accurate but less interpretable method. The existing methods for analysis of single-cell multimodal data vary in terms of explainability. Deep learning models tend to be less explainable. The field of explainable deep learning is rapidly developing, with many attempts at constructing explainable models [68]. However, learned parameter values of neural networks are often impossible to interpret. They represent meta properties of data, understandable by the computer, creating black-box models. New developments are needed for reliable large-scale single-cell applications. The explainability of a model can be increased with the use of interpretable parameters. For example, WNN produces cell-specific modality weights that inform which modality is most informative for each cell. At the same time, MOFA+ computes a distribution of modality features for each latent dimension of the reduced embedding. Applying such methods to multimodal single-cell data creates a joint low-dimensional embedding together with additional interpretable background information about cells or features, increasing the explainability of the output. Ocelli does increase explainability, integrating the advantages of the abovementioned methods to explore and visualize developmental multimodal single-cell data. In applying Ocelli, we preprocess data using probabilistic topic modeling that denoises the data while preserving its continuous structure. We adapt diffusion maps to a multimodal setting that allows us to include local and global information from each modality. Moreover, we directly include transition probabilities, e.g., from RNA velocity or Waddington-OT, when generating low-dimensional embeddings, which improves visualization of cell lineages and branching decisions. Lastly, we reconstruct highly sparse features using multimodal imputations, which allows us to recover feature-to-feature and cell-to-cell correlations.

Methods modeling RNA dynamics such as Velocyto [25] or scVelo [26] model the transcriptional content of cells and predict future states. Based on the relative kinetics of unspliced and spliced RNA, the methods compute a network of transition probabilities between cells and build a ”velocity” vector field that portrays cellular trajectories. The vector field was routinely visualized on the UMAP [45] or FLE [50] that were built without including cell transition probabilities. Recently, the VeloViz [69] method utilized cellular transitions to visualize unimodal single-cell data. VeloViz obtains each cell’s direction toward its predicted transcriptional state. Then, it calculates the proximity between all cell pairs by comparing their spatial direction to the predicted transcriptional one. Cells with the smallest proximities produce a nearest neighbor graph that is visualized. However, the drawback of VeloViz is that it requires coordinates of current and projected future transcription states for each cell, which limits a spectrum of applicable methods that produce transitions between cells, e.g., it is unable to leverage Optimal Transport inferred transition probabilities [27]. In addition, VeloViz applies to the visualization of scRNA-seq data only.

Multimodal Diffusion Maps implemented in Ocelli allow us to impute the genome-wide features’ activity in single cells and recover feature-to-feature relationships or modality-specific cell-to-cell correlations. Scalable implementation of the imputation algorithm enables real-time imputations of feature sets of interest for large multimodal datasets that contain a million cells. We envision the single-cell multimodal diffusion-based imputation can enhance the analysis of highly sparse features originating from chromatin state studies, e.g., profiling of chromatin accessibility, histone modifications, or DNA-bound proteins. Single-cell multimodal profiling technologies that leverage scCUT&TAG-based approaches [13–16] use Tn5 and/or Tn5-conjugated proteins (e.g. pA-Tn5 or nano-Tn5) to cut genomic DNA at open chromatin regions or close to sites of histone modifications. In the case of multimodal profiling of chromatin states, only one or two modalities can be captured for the same region of genomic DNA from the same cell. In the case of more modalities studied, multimodal imputations are inevitable to infer the chromatin states at single-nucleosome and single-cell resolution. Moreover, the proposed multimodal imputations can enhance our ability to identify distal elements, e.g., enhancer-promoter connections, on a genome-wide basis from single-cell profiling studies [70] and facilitate the building of single-cell enhancer databases [71]. One apparent shortcoming of diffusion-based imputation is that smoothing feature activity levels results in real numbers of the imputed levels instead of counts. The imputed levels include ”averaging” of multiple processes, both technical and biological, such as inefficient capturing of feature activity levels in single-cell sequencing technologies, as well as stochastic events during gene expression related to the dynamic switching of chromatin states at promoters and enhancers and RNA/protein production in bursts. The stochastic nature of the dynamics of chromatin states, transcription, and translation makes it impossible to infer genuine feature activity levels in single cells. Imputations have to involve the ”averaging” of multiple stochastic processes.

Overall, we have found that Ocelli employs powerful tools to study and visualize extensive developmental multimodal single-cell data to yield critical insights into the behavior of differentiating cells. We have shown, that Ocelli is a flexible diffusion-based method harnessing information from any number and type of modalities, along with prior knowledge of cell-to-cell transitions. It generates rich low-dimensional representation of continuous high-dimensional single- and mmultimodal data, which facilitates detection of relevant features from genome-wide measurements.

## Methods

### Basic notation

We use the following notation. Matrices and vectors are written in bold. Superscript refers to modality, and the subscript to coordinates. As a result, *ℳ* = **M**^1:*ℳ*^ represents multimodal data consisting of *M* modalities, where modality 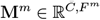comprises *C* cells and *F*^*m*^ features. 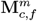 is an activity value of feature *f* from cell *c* in modality *m*.

### Multimodal weights

We model cell differentiation as a process of cellular movement in the latent space of single-cell features. According to such an interpretation, expanding distances in the cell’s neighborhood indicate a developmental process. Inspired by previous work on WNN [43], we quantify this expansion using multimodal weights, i.e., cell-specific distributions over modalities, assessing the likelihood of a developmental process occurring in modalities. The values of the weights are computed by comparing distances from a cell to its cross-modality nearest neighbors.

The computation of multimodal weights for cell *c* proceeds as follows. Let 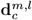be a vector of distances between cell *c* and its nearest neighbors from modality *l* in the feature space of modality *m*. Intuitively, 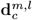encodes cross-modality behavior of cell *c* neighborhood from modality *l* in modality *m*. Then, we normalize cross-modality distances using modality-specific empirical cumulative distribution functions (ECDFs) of distances between cells. ECDF of modality *m*, denoted ECDF^*m*^, scales distances to a [0, 1] range according to the distribution of distances between cells in modality *m*. We estimate ECDFs by uniformly sampling *P* pairs of cell embeddings from each modality and calculating distances between them. The value of *P* depends on the total number of cells; however, usually, *P* = 1, 000 is enough to generate robust results. Scaled distances, denoted ECDF^*m*^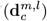, measure the expansion of the neighborhood of cell *c* from modality *l* in modality *m*, with respect to the global structure of modality *m*. Next, we score each modality as a sum over the remaining modalities of ECDF-scaled median distances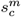is a score for modality *m* in cell *c*.

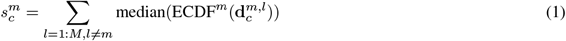

Then, we smooth the 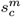 scores by averaging them over the nearest neighbors of cell *c* from the highest-scoring modality. Lastly, we normalize updated scores using a softmax function with the parameter *α*. The value of *α* should be greater or equal to 1; we used *α* = 10. Normalized scores produce cell-specific multimodal weights 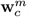 which sum to 1 for each cell.

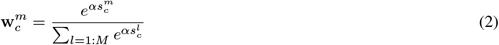

### Algorithm description

The algorithm is a multimodal generalization of the diffusion maps [20, 28]. The underlying idea is that high-affinity cells, i.e., cells close to each other in the single-cell feature space, are phenotypically closely related, and the direction in this space describes the differentiation trajectory. Diffusion maps model the resulting probabilistic diffusion process as a Markov chain, which is then eigendecomposed into a low-dimensional embedding. Multimodal diffusion maps (MDM) create a multimodal Markov chain that extends this approach to multimodal data.

MDM models multimodal data *ℳ* as a diffusion process. Firstly, MDM separately computes the affinities between cells and their nearest neighbors for each modality **M**. This step is performed using a density-adjusted symmetric Gaussian kernel *κ*

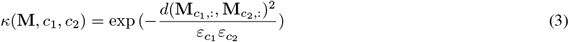

Equation (3) shows a formula for cell affinity *κ* between cells *c*_1_ and *c*_2_ in modality **M**. *d*(,) is the Euclidean metric and *ε*_*c*_ denotes a distance from cell *c* to its *N* ^th^ neighbor to account for local data density (usually *N* = 20 is enough for robust results). MDM models each modality as an unimodal Markov chain. The diffusion process of modality *m* is represented by a square matrix 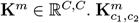 is a weighted affinity between cells *c*_1_ and *c*_2_.

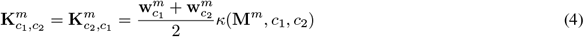

Next, MDM constructs the multimodal diffusion process **K** by element-wise summing weighted and row-normalized matrices **K**^1:*ℳ*^. The normalization is conducted using diagonal matrices **D**^*m*^ such that 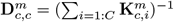

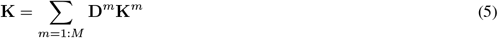

For efficient eigendecomposition to eigenvectors and eigenvalues, MDM converts **K** to a matrix 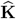 that has the following properties: (i) symmetric 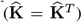, (ii) positive semi-definite 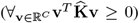, and (iii) non-negative 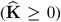. These properties guarantee to be a Hermitian matrix with real and non-negative eigenvalues. MDM row-normalizes (using a diagonal row-normalization matrix *D* such that 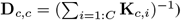 and then symmetrizes it along the diagonal to meet the conditions mentioned above.

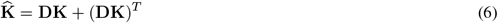

MDM eigendecomposes 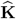 to create the low-dimensional embedding of multimodal data *ℳ* Eigendecomposition factorizes 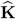 to a product of the eigenvector matrix **Q** (where **Q**_:,*i*_ is the *i*^th^ eigenvector), the eigenvalue vector **Λ** (where **Λ**_*i*_ is the *i*^th^ eigenvalue), and **Q**^*−* 1^.

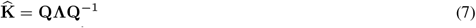

Let 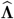 be the sorted eigenvalue vector with a decreasing order and 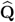 the eigenvector matrix with columns sorted accordingly. The *N* -dimensional MDM embedding of multimodal data *ℳ*, denoted MDM(*ℳ*), is defined as follows.

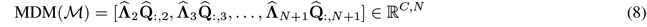

Ocelli employs scikit-learn [72], an ARPACK [73] wrapper, to compute the top *N* + *E* + 1 eigenvalues and corresponding eigenvectors. The quality of the computed eigenvectors may degrade as eigenvalues decrease; hence, *E* more than necessary eigenvectors are computed and discarded; we use *E* = 10. MDM ignores 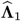, the largest eigenvalue, as it is non-informative in diffusion maps to recover developmental processes [20].

### Joint visualization of MDM components

MDM-generated low-dimensional embedding of multimodal data can be visualized in 2 or 3 dimensions using any dimension reduction algorithm. Ocelli has built-in wrappers for UMAP (umap-learn v0.5.3) [45] and ForceAtlas2 (v1.0.3) [50], which are recommended for well-clustered (e.g., postmitotic cells) or well-connected data (e.g., differentiating cells), respectively.

ForceAtlas2 is an algorithm for force-directed graph visualization. Ocelli provides three methods for converting MDM embeddings into graphs: (i) the nearest neighbors-based graph connects the nearest neighbors in the MDM latent space; (ii) the transitions-based graph connects a cell to its nearest neighbors in the MDM latent space with the highest developmental transition probabilities, e.g., from RNA velocity or Waddington-OT [25–27]; (iii) in the transitions-based graph, if additional information about cell timestamps is available, the nearest neighbors in the MDM latent space are selected only among cells from the subsequent timestamp (except cells from a terminal timestamp, when nearest neighbors are found in the same timestamp). MDM latent space captures information about the global structure of the data. Involving transitions between cells orders them locally in the visualization. A constructed graph has a predefined number of edges coming out of each node that is constant across all cells. If fewer cells have non-zero transitions in a cell’s MDM neighborhood, the graph edges link to the unconnected MDM nearest neighbors.

### Single-modality data exploration

Ocelli uses topic modeling to employ the multimodal analysis workflow to explore unimodal single-cell data. Ocelli splits data into latent single-cell modalities using latent Dirichlet allocation (LDA) [74], a probabilistic generative algorithm for learning relationships (called topics) between features. LDA models each cell as a multinomial mixture of topics, represented as distributions over features. Ocelli groups single-cell features according to their most probable topics and leaves only the top features according to LDA’s feature-topic distribution. The constructed groups form topic-based latent modalities.

Ocelli can visualize unimodal single-cell data by (i) constructing latent modalities using topic modeling and (ii) training MDM on latent modalities with LDA’s cell-topic multinomial distribution used as weights. This procedure reduces the inherent noise of single-cell data as we train MDM only on topic-specific features, and the impact of each latent modality is cell-specific using LDA’s cell-topic distribution for MDM weights.

### Diffusion-based multimodal imputation

Diffusion-based models can impute values of sparse single-cell features [19] by iteratively applying a diffusion operator to a feature count matrix **G**. For example, MDM produces a multimodal diffusion operator 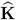. The *t*-step imputation proceeds as follows.

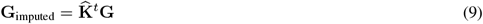

Eigendecomposition of 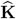 from equation (7) greatly simplifies running multiple diffusion steps.

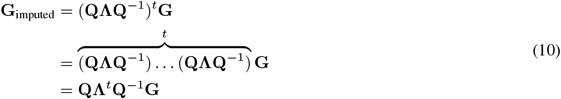

The only difficulty with equation (10), the computation of the inverse matrix *Q*^*−* 1^, can be evaded. 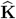 is symmetric, and ARPACK [73] produces unit-norm eigenvectors, making the eigenvector matrix **Q** orthonormal. Since the inverse of an orthonormal matrix equals its transpose, we derived an efficient imputation procedure with a constant complexity as the number of steps *t* varies.

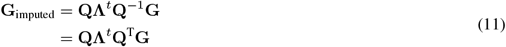

Ocelli imputes values of sparse multimodal single-cell features with an approximated version of the abovementioned procedure. It uses MDM-generated eigenvectors and eigenvalues in place of **Q** and **Λ**, respectively.

We investigated the computational time of Ocelli’s imputation on a 16-CPU machine in two experiments with *t* = 1 imputation steps (complexity is constant as *t* changes). (i) We fixed the number of features to 1,000 and generated sparse matrices with 10,000, 50,000, 100,000, 500,000, and 1,000,000 cells. (ii) We fixed the number of cells to 50,000 and generated sparse matrices with 1, 10, 100, 1,000, and 10,000 features. Values of matrices were sampled from a binomial distribution that samples 1 with probability *p* = 0.2 and 0 otherwise. We generated 50 eigenvalues and corresponding eigenvectors from a uniform distribution defined on a unit segment [0, 1].

### Benchmarking simulated data

We created and investigated two simulated developmental datasets. The binary tree dataset has three modalities and 6,000 observations grouped into six types (A-F). Modalities 1, 2, and 3 introduce developmental lineages A-B, C-D, and E-F. We sampled observations from 3-dimensional Gaussian distributions and then downsampled them to 1,000 observations per type with probabilities accounting for local densities. Equation (12) defines the distribution behind the downsampling procedure. *P* (*x*) is a probability of observation *x* being downsampled and NN_*N*_ (*x*) is the *N* ^th^ nearest neighbor of observation *x*. We used *N* = 25.

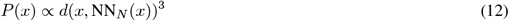

The developmental process was introduced to the dataset by assigning order to observations (pseudotime) based on the distance to manually selected developmental starting points. Then, we added noise by permuting every 100 cells along the pseudotime. During the multimodal analysis with Ocelli, we computed an MDM embedding (parameters: 10 dimensions, 20 nearest neighbors) and visualized it with ForceAtlas2 as a 30-edge-per-node nearest neighbor graph.

The rare transitions dataset has 4,500 observations grouped into nine types (A-I). Modalities 1 and 2 introduce developmental lineages A-C and D-I, respectively. Observations A-C in the first modality were sampled uniformly from 3-dimensional segments. The remaining observations originated from Gaussian distributions as in the binary tree dataset but resulted in 500 observations per type after downsampling. Pseudotime and noise were introduced analogically, as in the binary tree dataset. During the multimodal analysis with Ocelli, we computed an MDM embedding (parameters: 20 dimensions, 60 nearest neighbors) and visualized it with ForceAtlas2 as a 60-edge-per-node nearest neighbor graph.

We evaluated the quality of multimodal weights computed with MDM and WNN [43]. Weights from both methods were calculated using 60 nearest neighbors and compared to target weights using the mean squared error. Target modality weights equal 1 for lineages introduced in the modality and 0 for the remaining lineages.

### Preprocessing and analysis details of single-cell datasets

#### Hair follicle differentiation (SHARE-seq)

The hair follicle dataset is a subset of a SHARE-seq dataset [6] for female-mouse dorsal skin, containing 7,160 cells. We generated the RNA count matrix from BAM files using Velocyto v0.17.17 [25], producing the gene expression matrix with unspliced and spliced layers. We kept transit-amplifying cells (labeled *TAC-1, TAC-2*), inner root sheath cells (labeled *IRS*), and hair shaft cells (labeled *Medulla* and *Hair Shaft-cuticle*.*cortex*). When filtering the expression matrix, we performed the following steps: (i) removed Malat1, Gm42418, and AY036118 genes associated with RNA contamination; (ii) removed cell doublets using Scrublet v0.2.3 [75] with a predefined estimated collision rate [6];

(iii) removed cells with less than 50 expressed genes; (iv) removed genes with less than 50 counts; (v) removed genes highly correlated with the gene signature of cell proliferation [27]. The preprocessing steps have been carried out with Scanpy v1.9.1 [76]. In step (v), we computed Pearson correlations between the log-normalized expression of genes and the mean z-scores of the proliferation gene signature. Genes with a Pearson correlation larger than 0.18 were discarded. The resulting RNA count matrix had 7,160 cells and 6,731 genes. Then, we computed 50 LDA topics (scikit-learn v1.0.2 [72]) of the count matrix and scVelo’s v0.2.4 [26] stochastic RNA velocity transitions on 1,000 log-normalized highly variable genes.

We generated the ATAC-seq count matrix using Signac v1.10.0 [77] by mapping chromatin accessibility fragments to gene regions, including the 2kb upstream region. Next, we filtered out genes with less than 50 counts and genes correlated with the proliferation gene signature (the same genes as in RNA-seq). The resulting ATAC count matrix had 7,160 cells and 17,495 genes. Then, we computed 50 LDA topics of the count matrix.

During the multimodal analysis with Ocelli, we computed a low-dimensional MDM embedding trained on LDA topics (parameters: 20 dimensions, 20 nearest neighbors). We visualized it with ForceAtlas2 as a 3-edge-per-node nearest neighbor graph in 3 dimensions, which we then projected onto 2 dimensions. The gene signature for annotation of terminally differentiated cells were taken from SMART-seq2 Ana-VI hair follicle dataset [52]. Cuticle cell gene signature scores from IRS and HS [52] overlapped with Huxley’s and Cortex gene signature scores, respectively. The Cuticle cells in high-throughput scRNA-seq methods were indistinguishable from Cortex or IRs cells [6, 78] and were annotated jointly with them. For the sake of simplicity, we used only Huxley’s and Cortex annotation labels for cell identification.

#### Native hematopoietic differentiation (ASAP-seq)

The human bone marrow ASAP-seq dataset [12] contains 10,927 cells. We used matrices preprocessed by the dataset authors, namely, the protein count matrix with 238 protein tags and the ATAC-seq gene activity matrix with 3,000 genes (chromatin accessibility fragments were mapped to gene regions, including the 2kb upstream region, using Signac [77]).

#### Pancreatic endocrinogenesis (scRNA-seq)

The pancreatic endocrinogenesis RNA-seq dataset [65] contains 3,696 cells sampled from embryonic day 15.5. We used the published [26] gene expression matrix with unspliced and spliced layers. We filtered out cells expressed in less than 20 genes and removed genes expressed in less than 20 cells in both unspliced and spliced layers. The resulting count matrix has 3,696 cells and 5,465 genes. Next, we computed 20 LDA topics of the count matrix and generated topic-based modalities. Then, we calculated scVelo’s [26] stochastic RNA velocity transitions on 1,000 log-normalized highly variable genes.

#### Induced pluripotent stem cell reprogramming (scRNA-seq)

The induced pluripotent stem cell reprogramming scRNA-seq dataset [27] contains 259,155 cells. The RNA-seq count matrix was generated from BAM files using Velocyto [25]. When filtering the count matrix, we carried out the following steps: (i) kept cells from day 0 to 8 and cells in serum condition (from day 8.25 to 18); (ii) removed doublets using Scrublet [75] with a doublet score threshold set to 0.3; (iii) maintained cells from days 8-18; (iv) removed cells with less than 2,000 UMIs; (v) removed genes expressed in less than 50 cells; (vi) downsampled cells to 15,000 UMIs. Steps (iv)-(vi) followed the preprocessing instructions from the dataset authors. The resulting count matrix had 68,703 cells and 16,817 genes. Next, we computed 20 LDA topics of the count matrix and generated topic-based modalities.

## Benchmarking scalability

Scalability is a significant challenge for emerging computational methods in high-throughput single-cell genomics. We tested the scalability of Cobolt v0.0.1 [35], Matilda (first release version) [37], MDM, MOFA+ v0.7.0 [44], MOJITOO v1.0 [48], MultiVI v.0.19.0 [38], scMM v1.0.0 [36], and WNN (implemented in Seurat v4.9.9) [43] methods.

Benchmarking was conducted on matrices generated from the hair follicle dataset [6] in the following steps: (i) we sampled 10,000, 50,000, and 100,000 cells from the dataset, (ii) we selected 1,000 highly variable genes for RNA-seq modality, (iii) we selected 1,000 highly variable genes from gene activity matrix (generated using Signac [77] by mapping chromatin accessibility fragments to gene regions, including the region 2kb upstream from the promoter) and 1,000 highly variable peaks for ATAC-seq modality, (iii) we modified counts of each cell by randomly adding single counts to 10 of its features per modality to avoid cells with the same levels of features. We used a peak count matrix for methods that require or recommend peaks instead of gene activities. Data has been preprocessed using Scanpy [76]. We ran all methods on a 16-CPU machine, with an additional NVIDIA Tesla T4 GPU if the method requires GPU-based computation. We measured only the training time of each method, excluding data preprocessing and downstream visualization. We employed the following training procedures: (*Cobolt*) RNA-seq genes and ATAC-seq peaks with GPU trained for 100 epochs; (*Matilda*) RNA-seq genes and ATAC-seq gene activities with GPU trained for 100 epochs; (*MDM*) RNA-seq genes and ATAC-seq gene activities without GPU trained on exact (ENNs, computed with scikit-learn v1.0.2 [72]) and approximate (ANN, computed with NMSLIB v2.1.1 [79] nearest neighbors; (*MOFA+*) RNA-seq genes and ATAC-seq gene activities with GPU trained in accurate convergence mode; (*MOJITOO*) RNA-seq genes and ATAC-seq gene activities without GPU; (*MultiVI*) RNA-seq genes and ATAC-seq peaks with GPU trained for 100 epochs; (*scMM*) RNA-seq genes and ATAC-seq peaks with GPU trained for 100 epochs; (*WNN*) RNA-seq genes and ATAC-seq gene activities without GPU. We specified the dimensionality of the resulting latent space to 20 if applicable.

## Data preprocessing for benchmarking

This section describes generating embeddings for cohesiveness, RNA velocity vector field alignment, and RNA velocity-based terminal state separation benchmarks. We employed the following preprocessing procedures, following the recommendations of method authors. *Cobolt*: We trained 20-dimensional latent spaces for 200 epochs on (i) 1,000 highly variable RNA-seq genes and ATAC-seq peaks detected in more than 1 % of cells; (ii) all RNA-seq genes and ATAC-seq peaks detected in more than 1 % of cells. *Matilda*: We trained 20-dimensional latent spaces for 200 epochs on (i) 1,000 highly variable RNA-seq genes and 1,000 highly variable ATAC-seq gene activities, (ii) all RNA-seq genes, and all ATAC-seq gene activities. *MDM*: We trained 20-dimensional latent spaces on 50 LDA topics trained on all RNA-seq genes and 50 LDA topics trained on all ATAC-seq gene activities with (i) exact (ii) and approximate nearest neighbors. *MOFA+*: We trained 20-dimensional latent spaces in slow (most accurate) convergence mode on (i) 1,000 highly variable RNA-seq genes and 1,000 highly variable ATAC-seq gene activities; all RNA-seq genes and all ATAC-seq gene activities. *MOJITOO*: We trained latent spaces on (i) 50 PCs of log-normalized 1,000 highly variable RNA-seq genes and 50 PCs of log-normalized 1,000 highly variable ATAC-seq gene activities; (ii) 50 PCs of log-normalized all RNA-seq genes and 50 PCs of log-normalized all ATAC-seq gene activities. The dimensionality of latent spaces was computed automatically, resulting in (i) 40 and (ii) 41 dimensions. *MultiVI*: We trained 20-dimensional latent spaces for 200 epochs on (i) 1,000 highly variable RNA-seq genes and ATAC-seq peaks detected in more than 1 % of cells; (ii) all RNA-seq genes and ATAC-seq peaks detected in more than 1 % of cells. *scMM*: We trained 20-dimensional latent spaces for 200 epochs on (i) 1,000 highly variable RNA-seq genes and ATAC-seq peaks detected in more than 1 % of cells; (ii) all RNA-seq genes and ATAC-seq peaks detected in more than 1 % of cells. *WNN*: We trained multimodal graphs on (i) 50 PCs of log-normalized 1,000 highly variable RNA-seq genes and 50 PCs of log-normalized 1,000 highly variable ATAC-seq gene activities, (ii) 50 PCs of log-normalized all RNA-seq genes and 50 PCs of log-normalized all ATAC-seq gene activities. The resulting latent spaces and graphs were reduced to 2 and 3 dimensions with UMAP using 20 nearest neighbors and a minimum distance of 0.1. We additionally computed 2- and 3-dimensional ForceAtlas2 embeddings for MDM using a transitions-based graph with 100 nearest neighbors and 3 edges per node. We generated each embedding for 10 random seeds and saved median benchmark scores to negate any stochastic effects of dimension reduction algorithms on benchmark results.

### Benchmarking cohesiveness of signatures

RNA-seq signatures and ATAC-seq motifs (called together ”signatures” in this section) are a powerful aid in understanding the biology behind single-cell data. This section describes a benchmark evaluating the coherency of embedded cells with similar signatures in each modality. We define the activity of a signature in a cell as a mean z-score of gene expression for RNA-seq signatures or a motif variability score for ATAC-seq motifs. The cohesiveness benchmark is based on two criteria, as follows. (i) Close neighborhoods of highly signature-active cells should also be signature-active. We select 5 % of cells *G* with the highest activity and compute a score *Ψ*_c_ for each selected cell *c* defined as a mean signature activity score among its *n* = 3 nearest neighbors. (ii) Signature-active cells should be in a cohesive embedding region; disconnected high-activity clusters may indicate a disorganized embedding. We employ a density-based clustering algorithm DBSCAN [54] to establish a connective structure among the 5% of most active cells. DBSCAN creates a graph connecting cells within *ε* distance from each other. We set *ε* equal to the embedding’s estimated median distance between cells based on 1,000 random pairs of cells. We consider the median distance a fair threshold of proximity between cells.

The cohesiveness score of a signature for an embedding is calculated as follows. We compute a score Ψ_*G*_ for each connected component *G ∈ G* of the connectivity graph. A lower Ψ_*G*_ value indicates a better cohesiveness within *G*.

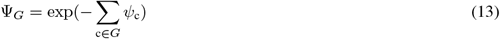

Then, we compute a total cohesiveness score Φ of an embedding over all connected graph components *G*.

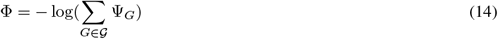

Φ penalizes graphs with multiple connected components. Note that the scale is inverted; higher values indicate a better-connected and locally more consistent signature portrayal.

We examined the cohesiveness scores on the hair follicle dataset [6]. For ATAC-seq, we used the 150 most-variable motifs computed using chromVAR v0.3 [80] from the peak count matrix. For RNA-seq, we ran a community-finding Infomap [81, 82] algorithm using the igraph v1.5.0 [83] package and kept clusters with more than 10 genes, resulting in 9 signatures. Cohesiveness scores were summed over all RNA-seq gene signatures for each evaluated embedding and divided by a maximum score so that the highest-scoring embedding scored 1. We analogically summed and scaled ATAC-seq motif cohesiveness scores. The final multimodal cohesiveness score is a sum of scaled RNA-seq and ATAC-seq scores with a maximum possible score of 2.

### Benchmarking RNA velocity vector field alignment

The RNA velocity alignment benchmark evaluates whether low-dimensional embeddings preserve RNA velocity dynamics. In the hair follicle dataset, we scored each cell as median cosine similarity between the cell’s projected developmental direction and vectors directed towards *n* = 3 cells with the highest RNA transition probabilities from the selected cell. Cosine similarities show high values if an embedding orders cells along the RNA velocity vector field. The score of an embedding is a median score across all cells. We computed projected developmental directions and RNA velocity transitions with scVelo [26] on log-normalized 1,000 highly variable genes.

### Benchmarking RNA velocity-based terminal state separation

The terminal state separation benchmark quantifies the separation of developmental lineage fates in low-dimensional embeddings. It uses absorption probabilities, which estimate how likely each cell is to transition into each identified terminal state based on RNA velocity. In the hair follicle dataset, we identified four terminal states and their absorption probabilities with CellRank v1.5.1 [53]. Let *T* be a set of terminal states. For each terminal state *t ∈ T*, we set its location as mean coordinates of 1 % of cells with the highest corresponding absorption probabilities. We score the separation of terminal state *t* as

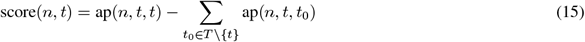

where ap(*n, t*_1_, *t*_2_) is a mean absorption probability of terminal state *t*_1_ among *n* nearest neighbors of the terminal state *t*_2_. score(*n, t*) rewards embeddings where absorption probabilities of terminal state *t* dominate its neighborhood. High absorption probabilities of remaining terminal states in *t*’s neighborhood penalize the score. We computed scores for *n ∈ {*20, 40, …, 980, 1000*}*.

## Code availability

Ocelli is an open-source resource, available under a BSD-Clause 2 license at https://github.com/TabakaLab/ocelli. The documentation and tutorials are available at https://ocelli.readthedocs.io.

## Data availability

Single-cell data are available at the Gene Expression Omnibus under accession numbers GSE140203 (hair follicle data), GSE156478 (human bone marrow data), GSE132188 (pancreatic endocrinogenesis data), GSE115943 (cell reprogramming data). Simulated data are available at https://doi.org/10.6084/m9.figshare.22700020.v1.

## Acknowledgments

We thank members of the Computational Genomics Group for their discussions. P.R. and M.T. are supported by the International Centre for Translational Eye Research (MAB/2019/12) project, which is carried out within the International Research Agendas Program of the Foundation for Polish Science, co-financed by the European Union under the European Regional Development Fund; and grants funded by National Science Center, Poland: the Sonata Bis 12 grant 2022/46/E/NZ2/0037 (M.T.), and Preludium 21 2022/45/N/NZ2/02311 (P.R.).

## Author Contributions

M.T. conceived and supervised the project. M.T. and P.R. designed and developed Ocelli. P.R. implemented Ocelli and wrote its documentation. M.T. and P.R. conducted the multimodal data analysis. M.T. and P.R. wrote the manuscript.

## Competing interests

M.T. and P.R. are named inventors on European Patent application EP23169340.9 relating to the work of this manuscript.

---

**Fig. S1.**
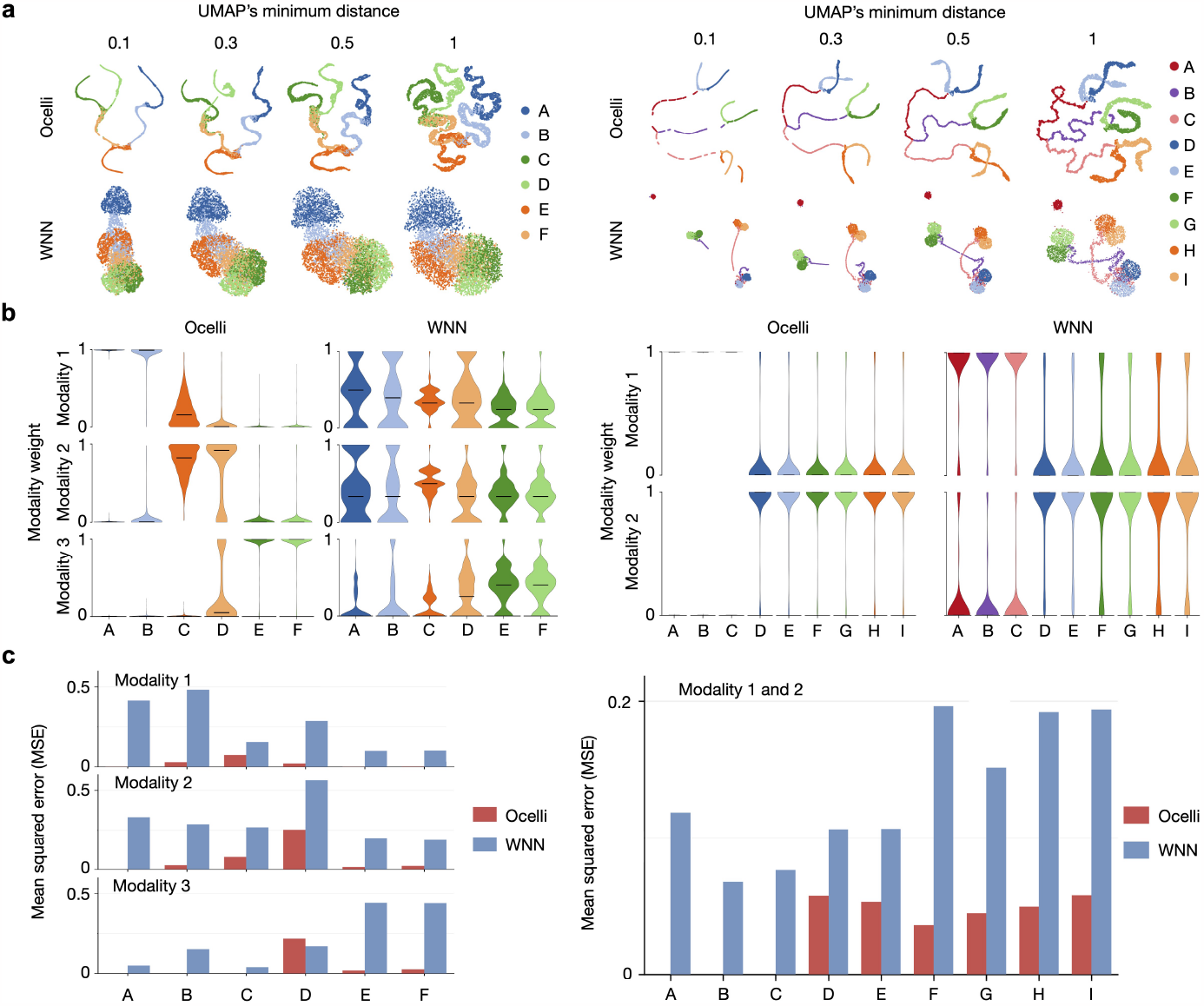
Reconstruction of developmental trajectories in simulated data with Ocelli (related to Fig. 1). Additional exploration of the binary tree dataset (left panel, see Fig. 1b) consisting of three modalities and six developmental lineages and the rare transitions dataset (right panel, see Fig. 1c) consisting of two modalities and nine lineages **a**, MDM and WNN embeddings visualized using UMAP with increasing values of the minimum distance parameter. **b**, Distributions of inferred multimodal weights computed using MDM and WNN. Violin plots show how informative modalities are for each cell lineage according to both methods. In the binary tree dataset, modalities 1, 2, and 3 introduce and are most informative about developmental lineages A-B, C-D, and E-F, respectively. In the rare transitions dataset, modalities 1 and 2 introduce and are most informative about developmental lineages A-C and D-I, respectively. **c**, Comparison of target and inferred weights computed using MDM and WNN. Target modality weights equal 1 for lineages introduced in the modality and 0 for the remaining lineages. The dissimilarity between inferred and target weights is calculated with the mean squared error (MSE). A lower error indicates a better sensitivity of weights.

**Fig. S2.**
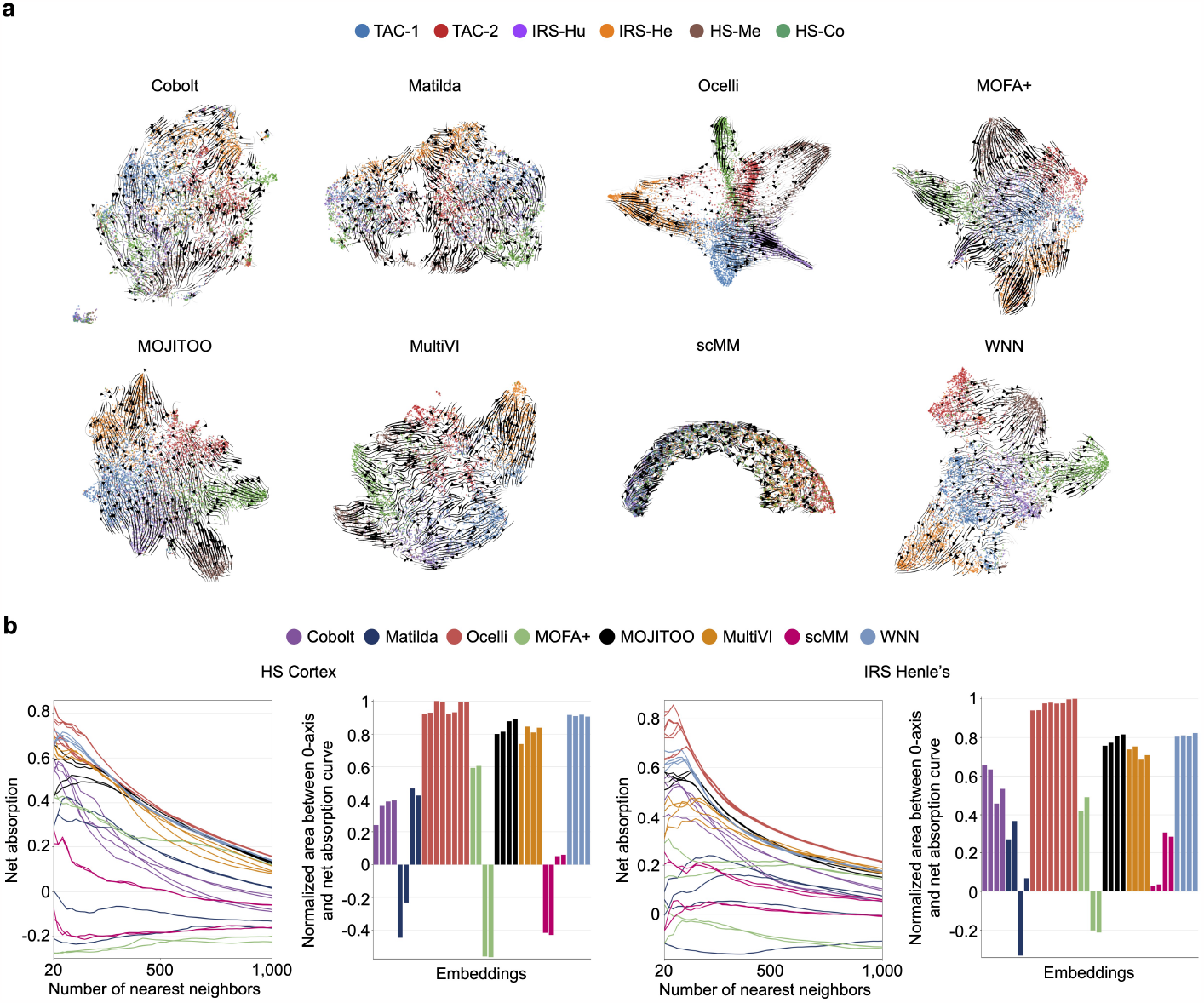
Additional analysis results from benchmarks (related to Fig. 3) **a**, Scatter plots show scVelo’s [26] RNA velocity projected onto 2D multimodal embeddings, generated from the analysis of hair follicle dataset by various methods. **b**, Net absorption scores of terminal states relating to hair shaft cortex and inner root sheath Henle’s layer cell types. Embeddings are ordered as in Fig. 3b.

